# Immigration and seasonal bottlenecks: High inbreeding despite high genetic diversity in an oscillating population of the biting midge, *Culicoides sonorensis* (Diptera: Ceratopogonidae)

**DOI:** 10.1101/2023.01.19.524819

**Authors:** Phillip Shults, Xinmi Zhang, Megan Moran, Lee W. Cohnstaedt, Alec C. Gerry, Edward L. Vargo, Pierre-Andre Eyer

## Abstract

Most population genetic studies concern spatial genetic differentiation, but far fewer aim at analyzing the temporal genetic changes that occur within populations. Vector species, including mosquitoes and biting midges, are often characterized by oscillating adult population densities, which may affect their dispersal, selection, and genetic diversity over time. Here, we used a population of *Culicoides sonorensis* from a single site in California to investigate short-term (intra-annual) and long-term (inter-annual) temporal variation in genetic diversity over a three year period. This biting midge species is the primary vector of several viruses affecting both wildlife and livestock, thus a better understanding of the population dynamics of this species can help inform epidemiological studies. We found no significant genetic differentiation between months or years, and no correlation between adult populations and the inbreeding coefficient (*F*_IS_). However, we show that repeated periods of low adult abundance during cooler winter months resulted in recurring bottleneck events. Interestingly, we also found a high number of private and rare alleles, which suggests both a large, stable population, as well as a constant influx of migrants from nearby populations. Overall, we showed that the high number of migrants maintains a high level of genetic diversity by introducing new alleles, while this increased diversity is counterbalanced by recurrent bottleneck events potentially purging unfit alleles each year. These results highlight the temporal influences on population structure and genetic diversity in *C. sonorensis* and provide insight into factors effecting genetic variation that may occur in other vector species with fluctuating populations.

## INTRODUCTION

The genetic structure of a population is influenced by many factors, such as dispersal, survival, adaptation, and population size. Through migration, selection, and genetic drift, these factors affect the amount and distribution of the genetic diversity within a population. While a population is considered a fundamental unit in evolution with implications in ecology, conservation biology, and population genetics (Hughes et al. 1997, Mimura et al. 2017), defining a population is not straightforward for some species. Within a single species, populations can be discrete or more continuous, but are often separated geographically, through seasonality, or by ecological associations (Waples and Gaggiotti 2006). When gene flow is high between two populations, their genetic structure can be quite similar, making them difficult to differentiate. To further confound this, the genetic structure of a population can change over time either through annual population phases or as a consequence of environmental changes (Jump et al. 2006). Temporal variations in genetic frequencies may occur at different rates and can have different outcomes between short-term (intra-annual) and long-term (inter-annual) change in genetic diversity (Fitzpatrick and Keller 2015).

Most genetic studies at the population level aim to investigate patterns of genetic structure occurring within and between geographically isolated populations (*i*.*e*., spatial genetic differentiation), but information on how genetic structure and diversity differ through time is often lacking (*i*.*e*., temporal genetic differentiation). Investigating temporal variation and the degree of genetic differentiation within a population can provide insights into the demographic stability of a population and its microevolutionary processes (Alonso-Blanco et al. 2005, Ayllon et al. 2006, Schwartz et al. 2007, Habel et al. 2014). Temporal genetic differentiation is influenced by the same factors as those affecting spatial genetic differentiation such as dispersal distance, immigration, selection, and population size (Loveless and Hamrick 1984, Waples and Gaggiotti 2006, Espíndola et al. 2012, Wogan and Wang 2018). For example, temporal variation in population size can influence genetic structure through a change in demographic processes between seasons resulting in recurrent or progressive bottlenecks. Smaller population sizes can result in greater inbreeding, changes in mating behavior or reproduction rate, and increased effects of genetic drift (Frankham 1996, Fraser 2008, Laikre et al. 2010). This scenario is exemplified in endangered populations, whereby reduction in the number of individuals over time gradually erodes genetic diversity (Hansen et al. 2012, Klinga et al. 2020). In contrast, increasing population size can enhance migration, increasing gene flow among populations and introducing new alleles from adjacent populations. Finally, changes in the environment can impact genetic diversity as different haplotypes or genotypes differentially survive or reproduce under distinct ecological conditions (Franks et al. 2007, Frachon et al. 2017). It is not uncommon for early and late season genotypes to arise, as environmental conditions vary by season favoring different alleles during distinct phases of a population cycle (*i*.*e*., temporally varying selection) (Borash et al. 1998, Hendry and Day 2005).

Populations can appear, disappear, expand, or contract with population size influenced by many factors that also affect the genetic diversity within a population. Populations with oscillating densities are predicted to exhibit lower overall genetic diversity than stable populations, due to high genetic drift occurring during periods of low density (*i*.*e*., recurrent bottleneck events) (Nei et al. 1975, Wright 1978, Whitlock 1992). Despite this prediction, many species maintain high genetic diversity within cycling populations (Ehrich and Jorde 2005, Porretta et al. 2007, Guardiola et al. 2016), owing to immigration during periods of increased population density (Matthysen 2005). In these instances, immigration introduces new alleles from surrounding populations allowing for rapid recovery of genetic diversity (Hansson et al. 2000, Keller et al. 2001, Ortego et al. 2007, Orantes et al. 2012). Disentangling how different factors impact genetic diversity is a major goal of population biology; and investigating patterns of population structure may lead to insights into evolutionary processes and the life-history traits affecting them.

A key question that has emerged from previous population genetic studies on vector insects, including *Culicoides* biting midges, relates to the role of fluctuating population size in shaping patterns of genetic diversity within populations. Biting midges are short-lived organisms producing multiple generations per year (Mullens 1987, Meiswinkel et al. 2014), with populations that oscillate annually between periods of high and low adult abundance (Gerry and Mullens 2000, Sanders et al. 2019). During times with low adult activity, it is assumed that genetic diversity is maintained in overwintering larvae (Campbell and Pelham-Clinton 1960, Mullens and Rodriguez 1992); however, some studies have suggested other means of surviving the winter months (Mullens and Lip 1987, Mayo et al. 2014). Regardless, the cyclical nature of these populations results in pronounced seasonal fluctuations in adult abundance and rates of migration. Most biting midge species studied so far exhibit long-distance dispersal, often wind-mediated (Ducheyne et al. 2007, Sanders and Carpenter 2014, Jacquet et al. 2016), leading to high gene flow among populations and an absence of population differentiation even at continental scales (Jacquet et al. 2015, Onyango et al. 2015a, Shults et al. 2021b). Despite the absence of genetic differentiation among most populations, high levels of inbreeding coefficient (*F*_*IS*_) and heterozygote deficiencies are commonly found in *Culicoides* species (Onyango et al. 2015b, Mignotte et al. 2021, Shults et al. 2021b). This is similarly reported for some mosquito species (Lehmann et al. 1997, Porretta et al. 2007, Fonseca et al. 2009, Goubert et al. 2016). This paradoxical finding may result from migration among populations followed by active mate choice (preferred mating with close relatives within populations), or from substantial seasonal variation in adult densities, frequently as observed for *Culicoides* species (Venter et al. 1997, Gerry and Mullens 2000, Meiswinkel et al. 2014) resulting in increased inbreeding and enhanced genetic drift when adult density is low. However, it should be noted that low abundance can also lead to sampling bias, whereby the few individuals collected at a single site may belong to the same cohort of closely related individuals due to synchronous emergence. In contrast, inbreeding in these species is expected to be reduced during high-density periods, through immigration and outcrossing with unrelated individuals. Considering these processes together, we predict that early season generations with less dense populations will have a high rate of inbreeding and low genetic diversity, while later generations with greater density will have increased genetic diversity.

In this study, we quantified the consequences of seasonal oscillation in adult population size on the amount and distribution of genetic diversity of the biting midge *Culicoides sonorensis* Wirth and Jones. This species is a confirmed vector of bluetongue virus, epizootic hemorrhagic disease virus, and vesicular stomatitis virus, and is a competent vector for a host of other pathogens (McGregor et al. 2022). We used monthly demographic and genetic monitoring over a 3-year period to infer whether variation in the adult population size was associated with changes in genetic composition. Specifically, we assessed whether periods of low densities are associated with lower genetic diversity and higher levels of inbreeding compared to periods of high densities. We also investigated if any alleles or genotypes were preferentially found in different periods of the density cycle and whether this population experienced any bottleneck events. In studying temporal variation within this population, we also gained important insights that will help inform the sampling efforts in future *Culicoides* population genetic studies.

## METHODS

### Specimen collection

Host-seeking adult female *C. sonorensis* were captured from a southern California dairy (San Bernadino County) every other week from April 2018 through April 2021 (Zhang 2022). Midges were captured over a 24-h period that started at 8:45 am (9:45 am during DST) using a CDC type miniature suction trap (Model 512, J.W. Hock), without a light, mounted at the opening of a rotating trap to sort midge capture into separate 80 min. collection periods. Carbon dioxide (CO_2_) was released from a compressed gas tank and delivered to the trap opening at a constant rate of 1,000 ml/min to mimic the respiration of a nearly grown Holstein heifer (Roberts 1972, Gerry et al. 2001). Midges attracted to the vicinity of the trap were pulled into a collection jar containing soapy water. Traps were collected the following morning. For each collection date, 15 midges were arbitrarily selected from one or more collection jars and placed into 95% ethanol for preservation. On the few dates when <15 midges were captured over the full 24-h period, all midges captured on that date were preserved in 95% ethanol. Samples were stored at −20°C until shipment to Texas A&M for genotyping as described below.

### DNA extractions and genotyping

A subsample of eight individuals was selected arbitrarily from the 15 midges preserved from each collection date. Genomic DNA was extracted from these individuals using a modified Gentra Puregene extraction protocol (Gentra Systems, Inc. Minneapolis, MN, USA). A total of 13 microsatellite markers (*C43, C45, C47, C65, C226, C230, C244, C508, C728, C838, C927, C1241*, and *C2085*) from Shults *et al*. (2022) (Shults et al. 2021a) were used in this study. The forward primer of each pair was 5′-fluorescently labeled with 6-NED, VIC, PET, or FAM. These markers were amplified in standard simplex PCR conditions using a Bio-Rad T100 thermal cycler (Bio-Rad, Pleasanton, CA, USA). Each PCR reaction contained 2.0 μl of DNA, 0.75 μM of a primer pair, 5.0 μl of 5× reaction buffer, 0.15 μl of Taq, and 16.35 μl of deionized water. The cycling conditions used for the amplification of microsatellite markers were 95 °C for 3 min, followed by 35 cycles of 95 °C for 1 min, 57 °C for 1.5 min, and 72 °C for 2 min, with a final extension at 72 °C for 5 min. An ABI 3500 capillary sequencer with a LIZ500 internal standard (Applied Biosystems, Foster City, CA, USA) was used to visualize PCR products. Alleles were scored using Geneious v.9.1 software (Biomatters, Auckland, New Zealand) (Kearse et al. 2012).

### Population genetic analyses

Allele numbers, allele frequencies, levels of heterozygosity, and *F*-statistics, were calculated for each microsatellite marker and year using the R package ‘genepop’ version 1.1.7 (Rousset et al. 2020). All subsequent population structure analyses were also performed using this R package. Deviations from Hardy–Weinberg equilibrium (HWE) were tested for each marker. The linkage disequilibrium (LD) for each pair of loci was tested across all collecting events using Fisher’s method. A maximum likelihood estimation of null allele frequency was performed using the EM algorithm and Brookfield’s (1996) methods. To assess whether the genetic composition of the population differed between time points, the pairwise genotypic differentiation between collecting dates was tested using the log-likelihood *G*-test with a Bonferroni correction. To determine whether the difference in genetic composition between two time points increased with the time separating them, genetic isolation by time (IBT) was estimated using the genetic differences between collecting events (*F*_ST_ /1-*F*_ST_) versus pairwise time differences (Log10). A Mantel test was performed to test for IBT. Time was coded continuously by collection date from 1 to 32.

To determine whether new alleles are frequently or even cyclically introduced into the population, we calculated the number of private alleles at each locus and compared the mean number of private alleles between the different months and years. Similarly, we compared the number of private alleles and differences in *F*_IS_ during periods of high (> 500 individuals) or low (< 500 individuals) adult population density, as well as between seasons with collection months categorized into early (Jan to Apr), mid (May to Aug) or late seasons (Sept to Dec). Tests for the significance of population density on inbreeding as well as differences in the mean number of private alleles per year were performed in JMP Pro v.16. NeEstimator v.2.1 was used to estimate the effective population size (*N*e) for each year of the study using both the linkage disequilibrium and heterozygote excess methods (Do et al. 2014).

To detect possible signs of genetic bottlenecks within the population, we used the software Bottleneck v1.2 (Piry et al. 1999). This software identifies recently bottlenecked populations by their increased level of heterozygosity relative to expected heterozygosity after a founder effect (Cornuet and Luikart 1996). We performed analyses using both the infinite alleles model (IAM) and the two-phased mutation model (TPM) with a Wilcoxon sign-rank test at 95% single-step mutations, and a variance among multiple steps of 16 based on 1000 iterations. Additionally, the occurrence of recent bottlenecks in the population were also determined by graphically investigating the distribution of allele frequencies (Luikart et al. 1998). Populations with large and stable sizes are characterized by an L-shaped distribution of allelic frequencies (*i*.*e*., many alleles with low frequencies). In contrast, bottlenecks are usually associated with the loss of rare alleles; they therefore induce a mode-shift distortion toward more alleles at intermediate frequencies.

## RESULTS

Over the 36 months of trapping, 33,674 *C. sonorensis* were collected from this population with an average of 981 midges captured per trap night (range = 0 to 4,955) (Fig. 1a). Of these, 255 individuals were genotyped from 32 of the trap months; 4 months were not used due to low collection numbers (*N* = 0, 0, 1, and 3). Across all samples, there was an average of 33 alleles per locus, and the expected (*H*_E_) and observed heterozygosity (*H*_O_) averaged across these loci ranged from 0.756 to 0.945 and from 0.264 to 0.820, respectively (Table 1). All loci were found to deviate significantly from HWE due to an apparent lack of heterozygotes, but no linkage disequilibrium was detected. The average multilocus inbreeding coefficient (*F*_IS_) was 0.337, though this varied considerably among loci ranging from 0.065 to 0.714 (Table 1). The number of individuals collected (a proxy for adult population density) was not correlated to *F*_IS_ (linear regression, *P* = 0.122) (Fig. 1b), nor was there any significant difference in *F*_IS_ during periods of high or low population density (Students’ *t*-test, *P* = 0.148) (Fig. 1c). The amount of genetic diversity and estimates of heterozygosity were similar whether the dataset was grouped and analyzed by month, year, density, or season. To test the effects of null alleles on the levels of homozygosity, the proportion of null alleles present for each locus were estimated. Only five of the 13 loci were found to have null allele frequencies below 8% (the cut off value considered for a low proportion of null alleles). However, of the remaining eight loci, four of these had low amounts of missing genotype data (< 2%), inconsistent with what would be expected from having a high proportion of null alleles (Table 1, Fig. S1). This suggests a negligible level of null alleles at most loci, and thus, unlikely to be the sole reason for the elevated levels of inbreeding.

**Table 1.** Locus information. Included for each locus are the names, total number of alleles, expected heterozygosity (*H*_e_), observed heterozygosity (*H*_o_), inbreeding coefficient (*F*_IS_), fixation index (*F*_ST_), percentage of missing genotype, and estimate null allele frequency (Brookfield’s 1997 method). The bottom row is the multilocus average across all markers. For both the missing genotype and null allele frequency data, 8.0% and 0.08 (respectively) represent the cutoff values for low frequencies.

| Locus | Number of<br>alleles | $H_e$ | $H_o$ | $F_{IS}$ | $F_{ST}$ | Missing<br>genotypes (%) | Null allele<br>frequency |
| --- | --- | --- | --- | --- | --- | --- | --- |
| C43 | 23 | 0.756 | 0.667 | 0.119 | 0.0051 | 3.15 | 0.0481 |
| C45 | 28 | 0.920 | 0.674 | 0.268 | 0.0128 | 7.48 | 0.1255 |
| C47 | 33 | 0.874 | 0.817 | 0.065 | 0.0173 | 0.00 | 0.0272 |
| C65 | 22 | 0.843 | 0.772 | 0.084 | -0.0021 | 0.00 | 0.0357 |
| C226 | 50 | 0.945 | 0.310 | 0.672 | -0.0012 | 1.57 | 0.3244 |
| C230 | 39 | 0.920 | 0.570 | 0.380 | -0.0013 | 0.00 | 0.1797 |
| C244 | 22 | 0.886 | 0.448 | 0.495 | -0.0122 | 1.57 | 0.2319 |
| C508 | 26 | 0.921 | 0.264 | 0.714 | 0.0104 | 16.54 | 0.3426 |
| C728 | 31 | 0.923 | 0.820 | 0.112 | -0.0023 | 5.12 | 0.0506 |
| C838 | 47 | 0.875 | 0.667 | 0.238 | 0.0023 | 6.69 | 0.1083 |
| C927 | 20 | 0.829 | 0.321 | 0.613 | 0.0292 | 14.96 | 0.3092 |
| C1241 | 36 | 0.873 | 0.490 | 0.439 | 0.0038 | 1.57 | 0.2018 |
| C2085 | 55 | 0.892 | 0.779 | 0.127 | 0.0126 | 0.39 | 0.0573 |
| Multilocus<br>average | 33.23 | 0.881 | 0.584 | 0.337 | 0.0053 | 4.54 | 0.1571 |

**Figure 1.**
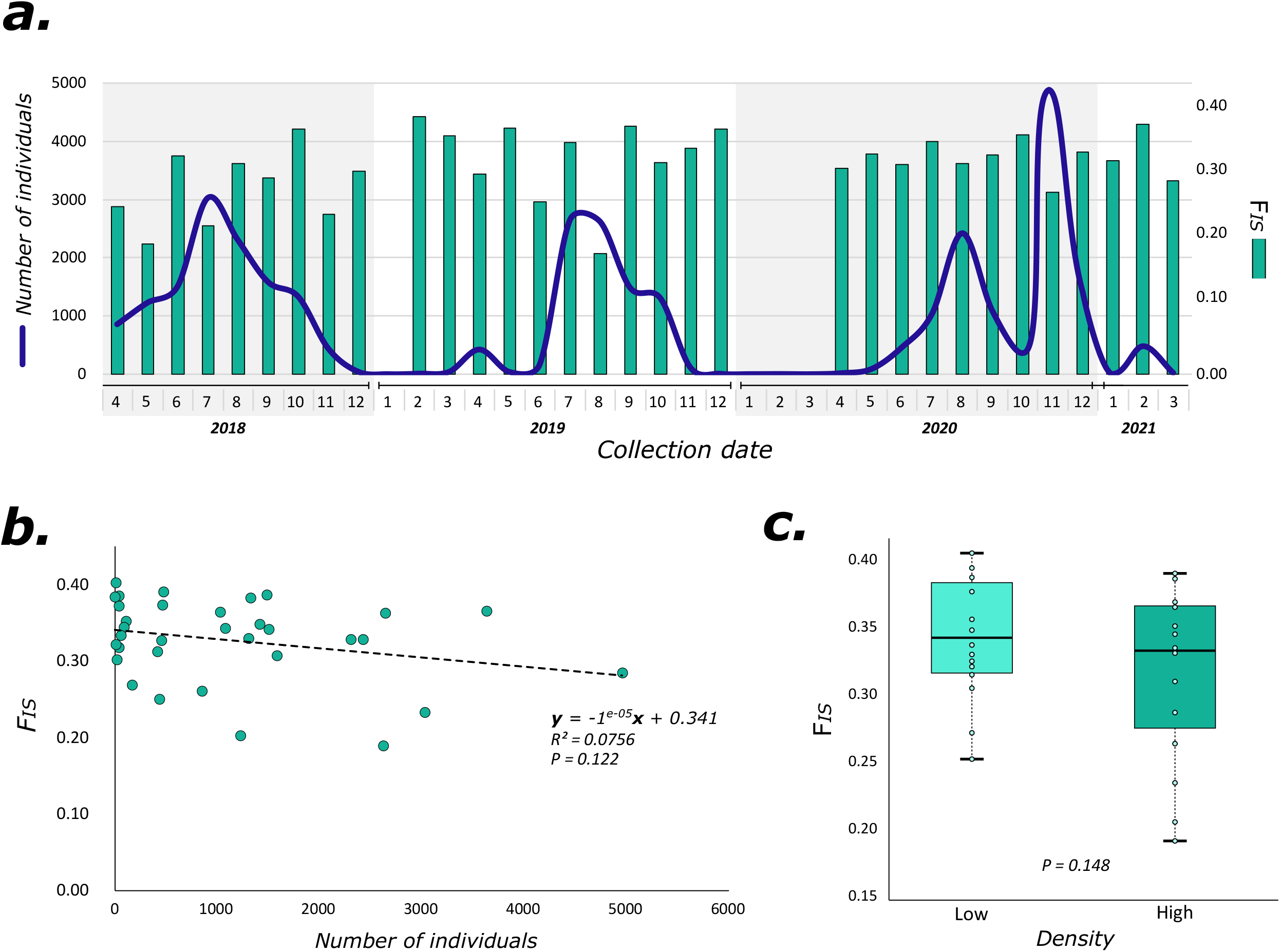
Relationship between population size and the inbreeding coefficient (*F*_IS_). (a.) The line shows the total number of individuals collected for each month and the bar shows the corresponding F_IS_. Months with no bars were not analyzed due to low collection numbers. (b.) The slightly negative, but nonsignificant (*P* = 0.122), correlation between population size and *F*_IS_. (c.) Comparison of the mean F_IS_ during periods of low and high population densities.

There was little to no genetic differentiation among months with an overall mean *F*_ST_ value of 0.005 and no isolation by time was present (Mantel test, *P* = 0.405) (Fig. 2). The log-likelihood *G*-test found midges collected on only two dates to be significantly different from each other (*P* = < 0.001; May 2018 and August 2020) out of a total of 528 comparisons. Similarly, there was no genetic differentiation between years with a pairwise *F*_ST_ close to zero (−0.00015). There were changes in allelic frequencies from year to year in 7 of the 13 loci as well as month to month in 11 of 13 loci (exact *G*-test, *P* ≤ 0.05); however, these changes in allelic frequency do not appear to be linked to any seasonal effect or adult population abundance. The overall frequencies remained relatively stable over time which indicates a sufficiently large effective population size. This was further supported by the *N*e estimates for each year which were determined to be infinitely large under both the linkage disequilibrium and heterozygote excess methods. The mean number of private alleles per locus was 4.80 from year 1 (Apr. 2018 – Mar. 2019), 3.10 from year 2 (Apr. 2019 – Mar. 2020), and 3.75 from year 3 (Apr. 2020 – Mar. 2021). There was no significant difference in the mean number of private alleles across years (ANOVA, *P* = 0.240), seasons (ANOVA, *P* = 0.238), months (ANOVA, *P* = 0.936) (Fig. 3a-c), or in relation to the number of adults collected (Students’ *t*-test, *P*= 0.878). However, each year, approximately 46 new alleles were introduced into this population, accounting for roughly 15% of the total number of alleles. While the mean number of private alleles was not significantly different among the seasons, nearly half of all private alleles were detected during the last four months of the year (Fig. 3c).

**Figure 2.**
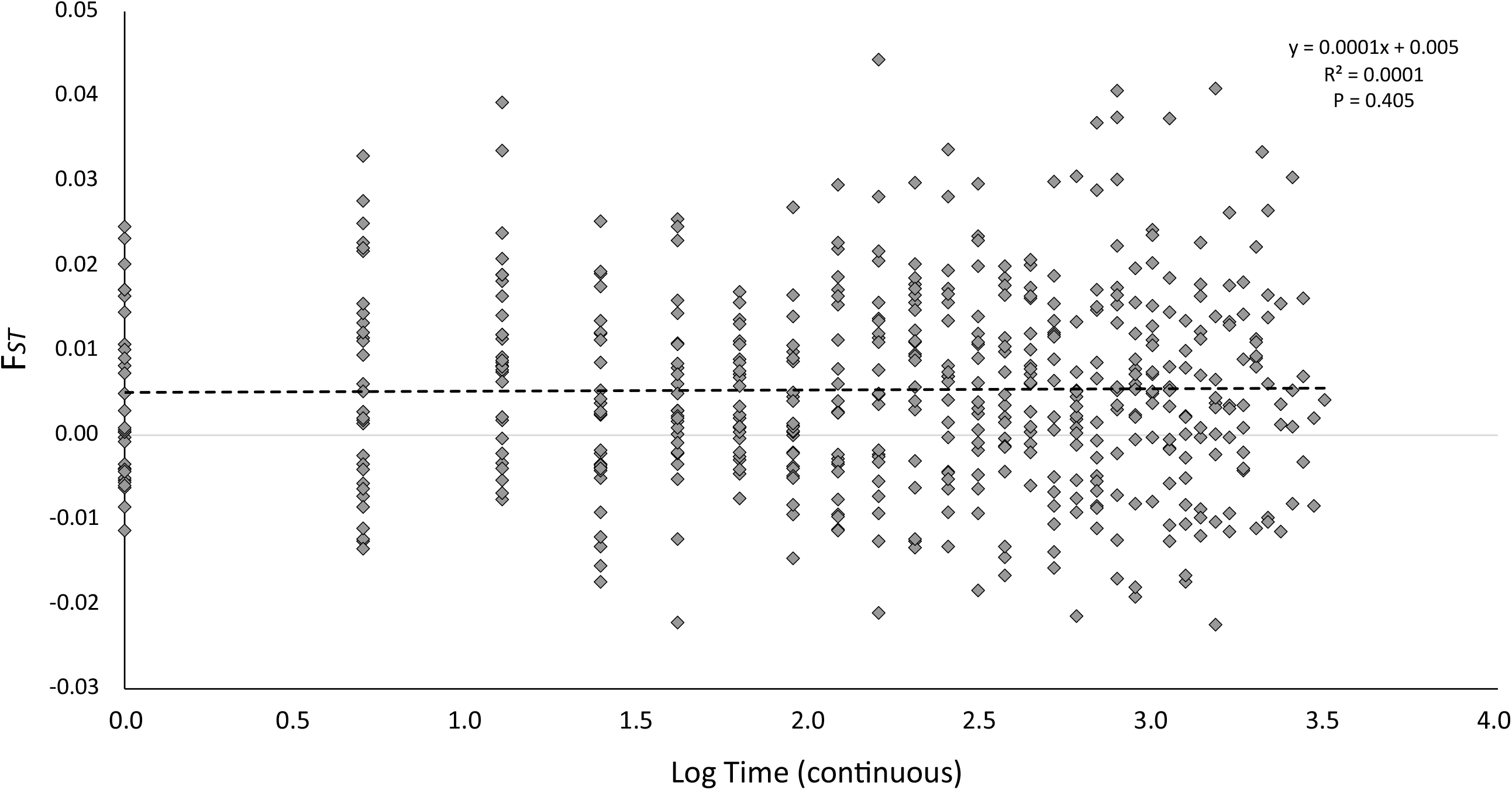
Isolation by time. The fixation index (*F*_ST_) plotted against the log transformed time. Time was coded continuously from 1 to 32, starting with April 2018 (1) and ending with March 2021 (32).

**Figure 3.**
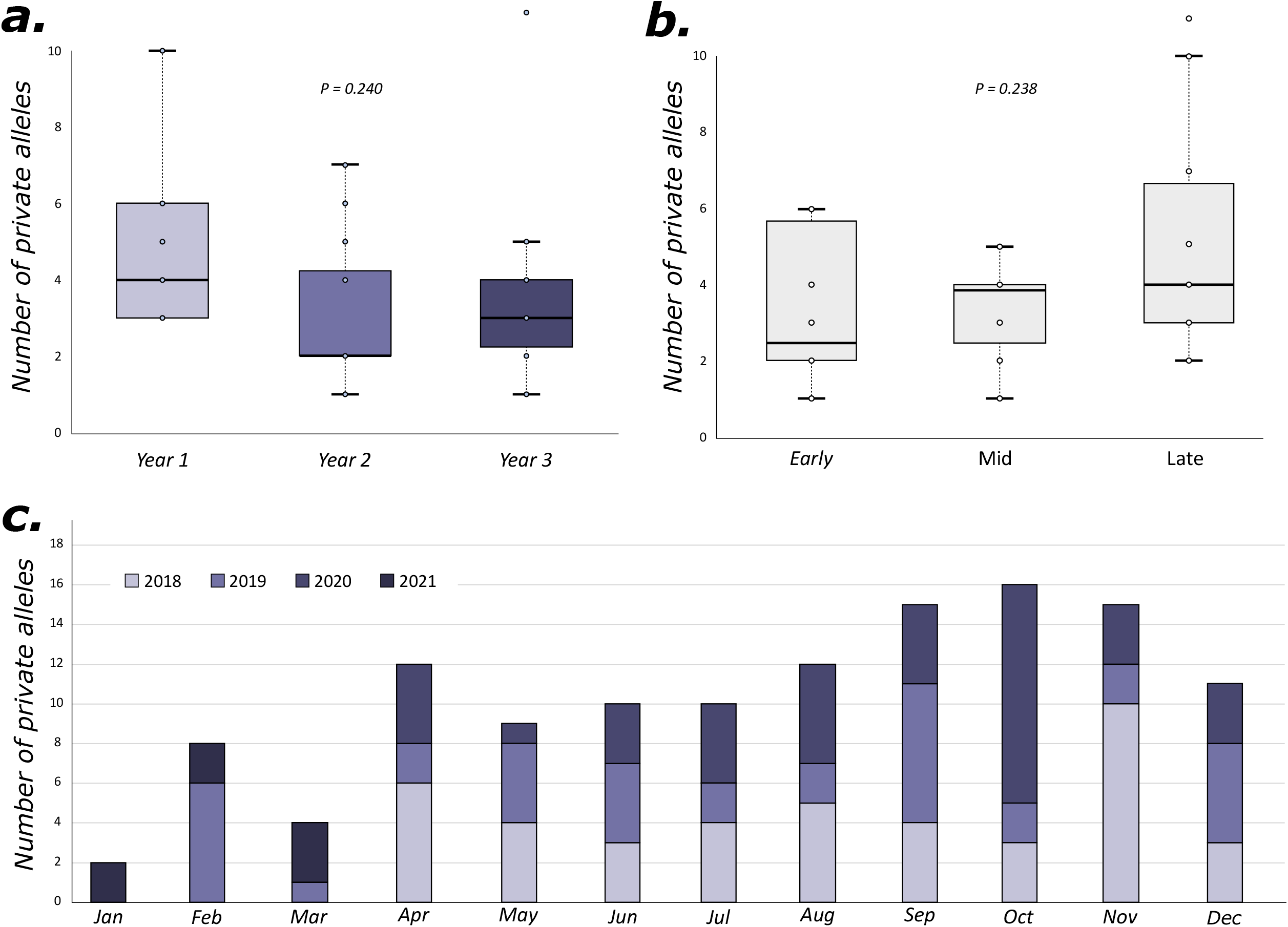
Private alleles withing the population. (a.) Comparison of the mean number of private alleles found in each year and (b.) during different seasons across all years. (c.) The total number of private alleles found in each month across all study years.

When testing for the occurrence of bottlenecks in each month separately, nine of the 32 months were found to have an excess of heterozygotes (*i*.*e*., a bottleneck event) under the IAM (Table 2, Fig. S2). Interestingly, significantly more bottleneck events were identified in this population during months of low adult abundance (*N* = 8) compared to months with a high adult population density (*N* = 1) (Likelihood ratio, *P* = 0.004). When testing for the occurrence of bottlenecks for each year, all were found to be in mutation-drift equilibrium under the IAM, suggesting that these intermittent bottlenecks are not enough to affect the overall genetic diversity (Table 2). This is further supported by the L-shaped distribution of allelic frequencies for all microsatellite markers, with an abundance of low-frequency alleles (0.001 to 0.1; Fig. 4a). The absence of a significant loss of rare alleles suggests a stable population; however, the fact that these are not the same alleles over time hint that at least some are being introduced. Finally, rarefaction was used to determine the allelic richness given N individuals. Approximately 50% of the total number of alleles were found in the first 27 individuals, and 75% and 95% of the total alleles were obtained from sampling 73 and 210 individuals respectively (Fig. 4b).

**Table 2.** Collection data, genetic diversity, and bottleneck information for the *C. sonorensis* used in study. For each collection date are the number of individuals collected and whether this was categorized as a high or low population density (cutoff value < 500 individuals); expected heterozygosity (*H*_e_), observed heterozygosity (*H*_o_), and inbreeding coefficient (*F*_IS_); and if a bottleneck event is inferred (*P* < 0.05) under the infinite allele model (IAM) and the two-phased mutation model (TPM). Collection dates in bold indicate a bottleneck event under the IAM.

| Collection Date | Individuals | Density | $H_e$ | $H_o$ | $F_{is}$ | IAM | TPM |
| --- | --- | --- | --- | --- | --- | --- | --- |
| 4/23/2018 | 857 | high | 0.850 | 0.615 | 0.261 | 0.3677 | 0.8633 |
| 5/23/2018 | 1227 | high | 0.854 | 0.684 | 0.203 | 0.5270 | 0.9045 |
| 6/18/2018 | 1507 | high | 0.851 | 0.566 | 0.342 | 0.0549 | 0.8633 |
| 7/30/2018 | 3028 | high | 0.855 | 0.662 | 0.232 | 0.1367 | 0.8121 |
| 8/29/2018 | 2304 | high | 0.855 | 0.584 | 0.328 | 0.2274 | 0.8121 |
| 9/26/2018 | 1583 | high | 0.848 | 0.587 | 0.307 | 0.2072 | 0.8302 |
| **10/24/2018** | **1331** | **high** | 0.856 | 0.532 | 0.383 | **0.0043** | 0.3177 |
| 11/21/2018 | 437 | low | 0.874 | 0.663 | 0.250 | 0.1527 | 0.7291 |
| 12/18/2018 | 39 | low | 0.888 | 0.619 | 0.318 | 0.2072 | 0.3177 |
| 1/21/2019 | 0 | - | - | - | - | - | - |
| **2/26/2019** | **9** | **low** | 0.861 | 0.526 | 0.402 | **0.0239** | 0.2709 |
| **3/26/2019** | **40** | **low** | 0.880 | 0.572 | 0.373 | **0.0020** | **0.0239** |
| **4/24/2019** | **423** | **low** | 0.876 | 0.615 | 0.312 | **0.0471** | 0.5270 |
| **5/24/2019** | **38** | **low** | 0.832 | 0.518 | 0.385 | **0.0341** | 0.6576 |
| 6/20/2019 | 167 | low | 0.860 | 0.637 | 0.269 | 0.2274 | 0.8473 |
| 7/15/2019 | 2647 | high | 0.856 | 0.556 | 0.362 | 0.0955 | 0.7061 |
| 8/13/2019 | 2621 | high | 0.865 | 0.713 | 0.189 | 0.4730 | 0.7291 |
| 9/9/2019 | 1482 | high | 0.833 | 0.534 | 0.387 | 0.1527 | 0.5803 |
| 10/9/2019 | 1304 | high | 0.857 | 0.583 | 0.330 | 0.1698 | 0.7061 |
| 11/6/2019 | 109 | low | 0.862 | 0.567 | 0.353 | 0.0636 | 0.4730 |
| **12/18/2019** | **7** | **low** | 0.865 | 0.551 | 0.383 | **0.0107** | 0.0549 |
| 1/13/2020 | 1 | - | - | - | - | - | - |
| 2/13/2020 | 3 | - | - | - | - | - | - |
| 3/27/2020 | 0 | - | - | - | - | - | - |
| 4/21/2020 | 17 | low | 0.814 | 0.560 | 0.322 | 0.1879 | 0.6576 |
| 5/19/2020 | 90 | low | 0.809 | 0.544 | 0.345 | 0.5270 | 0.8121 |
| 6/16/2020 | 460 | low | 0.853 | 0.585 | 0.327 | 0.1527 | 0.7291 |
| 7/14/2020 | 1036 | high | 0.855 | 0.560 | 0.364 | 0.1527 | 0.4197 |
| 8/12/2020 | 2422 | high | 0.808 | 0.545 | 0.329 | 0.2274 | 0.8121 |
| 9/8/2020 | 1085 | high | 0.866 | 0.580 | 0.342 | 0.1527 | 0.6823 |
| **10/6/2020** | **468** | **low** | 0.879 | 0.565 | 0.374 | **0.0199** | 0.1082 |
| 11/5/2020 | 4955 | high | 0.858 | 0.624 | 0.284 | 0.1879 | 0.6066 |
| 12/1/2020 | 1414 | high | 0.828 | 0.549 | 0.348 | 0.1219 | 0.5000 |
| 12/29/2020* | 61 | low | 0.871 | 0.579 | 0.334 | 0.0732 | 0.3424 |
| **2/23/2021** | **480** | **low** | 0.883 | 0.545 | 0.391 | **0.0009** | 0.2709 |
| **3/22/2021** | **22** | **low** | 0.863 | 0.611 | 0.302 | **0.0043** | 0.1698 |
| Year 1 | 12362 | - | 0.861 | 0.601 | 0.309 | 0.6086 | 0.9999 |
| Year 2 | 8802 | - | 0.856 | 0.586 | 0.330 | 0.1411 | 0.9999 |
| Year 3 | 12510 | - | 0.849 | 0.571 | 0.338 | 0.3619 | 1.0000 |

\* December 29, 2020 was used as the January 2021 collection for all analyses.

**Figure 4.**
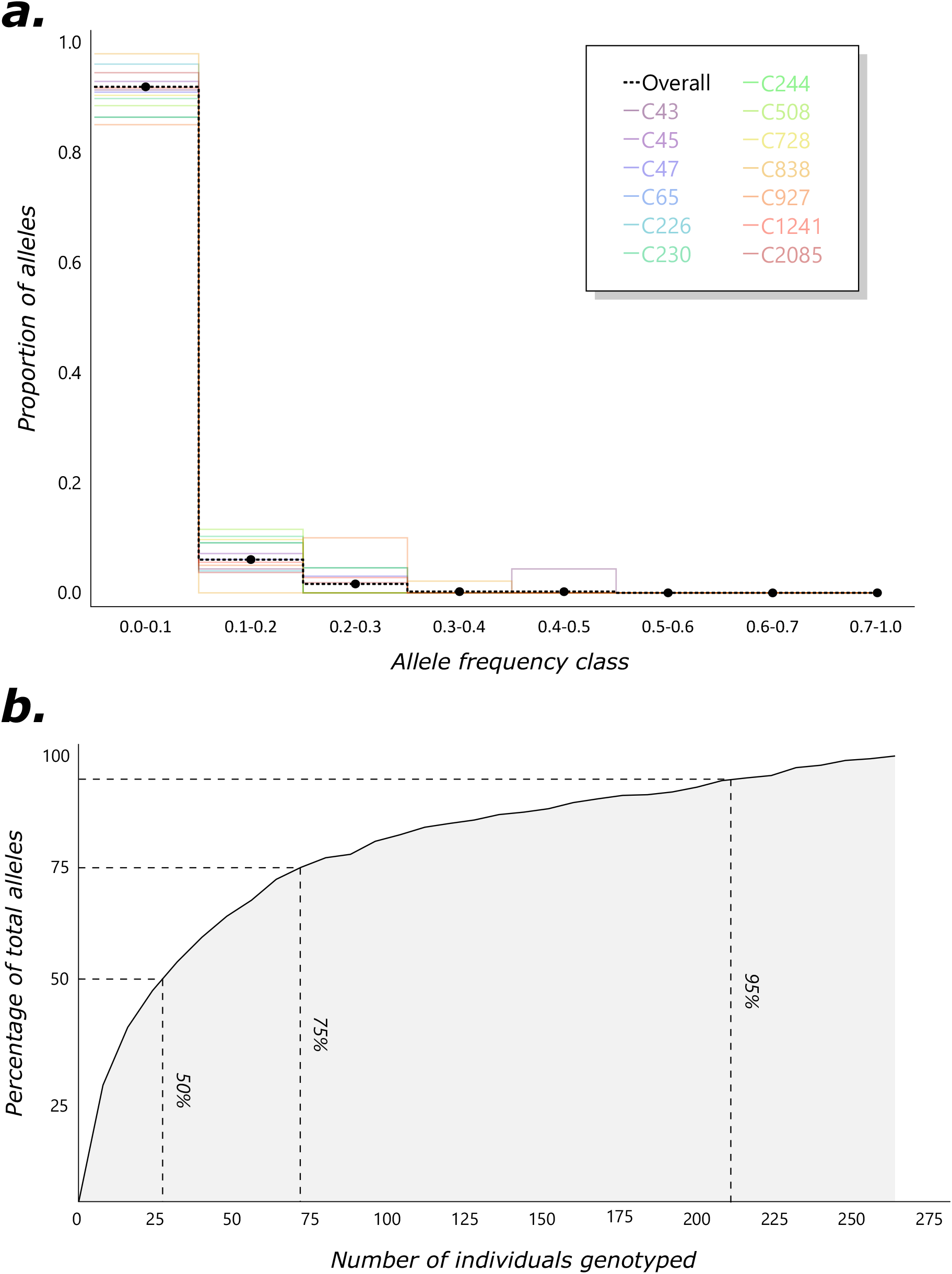
Allelic richness. (a.) The proportion of alleles that occur at various allele frequencies overall, and for each locus. (b.) The proportion of the total alleles as a function of the number of individuals sampled.

## DISCUSSION

Despite huge fluctuations in the adult population size, almost no genetic differentiation was observed. Cohorts of individuals collected at the beginning of the study were genetically indistinguishable to those collected three years later (Fig. 2). The mean *F*_ST_ value across all collection dates was near-zero indicating the genetic structure and genetic diversity of this population remained relatively constant across a multitude of generations (approx. 27-33) (Mullens and Lip 1987). This stability likely stems from the population’s capacity to support an incredibly large number of individuals, though certain patterns emerged during periods of high and low adult abundance. As the adult population density increased, new alleles were introduced through migration, followed by recurrent bottleneck events during periods with lower adult activity. These two forces appear to be counterbalancing each other while still allowing for the maintenance of a high level of genetic diversity. Additionally, despite having a rather large population, surprisingly few individuals were needed to determine the heterozygosity within this population. The mean levels of heterozygosity observed for each month (*N* = 8) were similar to the levels observed when analyzing entire years (*N* = 72 – 87) (Table 2). However, to obtain at least 50% of the overall allelic diversity, genotyping of approximately 25 individuals was required (Fig. 4b). There were also diminishing returns on the increase in allelic diversity gained when sampling more than 75 individuals, though it is likely that with immigration constantly bringing new alleles, it is impossible to sample all of the genetic diversity.

Immigration likely accounts for a large part of the genetic diversity found within the *C. sonorensis* population studied here as each locus harbored a high amount of rare and private alleles. Estimating migration using private alleles is more likely to reflect recent immigration as these alleles are assumed to be newly introduced in the population. While new alleles can arise via mutation, this occurs at a much slower rate than could account for the approximately 15% new alleles observed each year. The high number of rare and private alleles indicates that this population constantly receives genetic diversity from adjacent populations. Bottlenecks and genetic drift usually lead to random loss of alleles in sparse and relatively isolated populations (Frankham et al. 2002), but this was not observed in this biting midge population. While bottleneck events were inferred each year, these immigrants quickly replenished the genetic diversity within the population. From our data alone, it is not possible to determine the source of these introduced alleles; however, midge movement from 5-30 km is relatively common (Hendrickx et al. 2008, Sanders and Carpenter 2014) but the potential introduction from more distant populations is also possible (Ducheyne et al. 2007).

Like several species of mosquitoes, *Culicoides* populations have been described with remarkably high levels of inbreeding (Onyango et al. 2015b, Mignotte et al. 2021, Shults et al. 2021b). Our results also uncovered high *F*_IS_ values within this population, but contrary to our hypothesis, the level of inbreeding does not seem to originate from a perceived reduction in population size during periods of low adult abundance (Fig. 1). It should be noted that the number of adults collected in our traps may not reflect the true population size as there may be no reduction in the larval densities. However, for at least *C. sonorensis*, there is evidence that immature abundance also decreases during the winter months (Mullens and Lip 1987, Mayo et al. 2014), and further evidence from this population suggests continuous immature development and emergence (Zhang and Gerry 2023). Regardless, our results suggests that factors other than adult abundance are responsible for the elevated levels of *F*_IS_ commonly associated with populations of *Culicoides* biting midges. The first, and potentially simplest, being that these populations are in fact inbred; however, there are several other explanations for increased estimates of homozygosity that warrant further discussion.

Most biting midge species studied thus far, including *C. sonorensis*, are associated with an absence of genetic differentiation between geographically distant populations (Jacquet et al. 2015, Onyango et al. 2015a, Jacquet et al. 2016, Shults et al. 2021b), which reduces the possibility that the elevated inbreeding coefficient resulted from the Wahlund effect (*i*.*e*., sub-structuring within a population). Alternatively, the high level of inbreeding has often been attributed to the presence of a high number of microsatellite null alleles (*i*.*e*., an allele that consistently fails to amplify using PCR), as these can cause significant heterozygote deficits relative to HWE (Onyango et al. 2015a, Mignotte et al. 2021). However, the various methods developed to estimate null allele frequencies do so by testing for HWE and assume that the observed heterozygote deficiencies have no other origin (Dempster et al. 1977, Chakraborty et al. 1992, Brookfield 1996, Chapuis and Estoup 2007). These methods will therefore invariably suggest high frequencies of null alleles in every inbred population. In addition to artificially raising the level of homozygosity, the presence of null alleles also increases the amount of missing data, as individuals homozygous for a null allele will fail to amplify (Kalinowski and Taper 2006). Here, four of the eight loci identified as having a high proportion of null alleles in our study had less than 2% missing data, meaning that these methods are overestimating the presence of null alleles (Fig. S1). Additionally, the presence of null alleles is locus-specific and therefore unlikely to explain concordant heterozygote deficits across all loci investigated in our study (Chakraborty et al. 1992, Dakin and Avise 2004), as well as heterozygote deficits uncovered using SNP datasets (Shults et al. 2021b). Instead, we propose that the observed high inbreeding coefficient is being slightly inflated by the numerous alleles brought to this population by migrants (estimated at 46 per year). High estimates of expected heterozygosity would be inferred from populations with high numbers of migrants, as these individuals will carry newly introduced alleles. As inbreeding is estimated as a function of the expected and observed levels of heterozygosity, the elevated level of inbreeding in our study may stem from an inflated estimate of expected heterozygosity due to migration. In summary, we believe that this *C. sonorensis* populations is experiencing moderate levels of inbreeding but acknowledge that these estimates may be inflated to some degree.

This region of California has been very active in dairy production for many years (Gerry and Mullens 2000), and as such, there is a possibility that this population favors genes beneficial to utilizing artificial habitats. The overall genetic differentiation between *C. sonorensis* populations across the entire US is relatively low (Shults et al. 2021b), and an adaptation to livestock breeding habitats could help explain the range-wide genetic homogeneity and elevated inbreeding. Though inbreeding is generally associated with low genetic diversity, this is not what was observed in this population. The constant introduction of new alleles could be the mechanism by which this species overcomes inbreeding depression. Conversely, *Culicoides* populations might not suffer from inbreeding depression at all due to having extremely large populations (Bataillon and Kirkpatrick 2000). Inbreeding can even be beneficial in some population as it can quickly purge deleterious mutations by exposing them to selection through increase homozygosity (Barrett and Charlesworth 1991, Edmands 2007, Pujol et al. 2009). Potentially this, combined with the recurrent bottleneck events, could be counterbalancing the influx of new genetic variants each year by purging deleterious alleles; keeping genetic diversity high while still favoring local adaptation (Kirkpatrick and Jarne 2000). This annual cycle appears to be aiding in the maintenance of genetic diversity and affords this, and likely other *Culicoides* populations, the ability to rapidly adapt should environmental conditions change.

## ACKNOWLEDGMENTS

The authors would like to thank the dairy facility for allowing us access to collect biting midges. We would also like to thank the ABADRU unit of the USDA-ARS for support during manuscript preparation. This research was funded by the Urban Entomology Endowment fund from Texas A&M University, a USDA Non-Assistance Cooperative Agreement: 58-3020-9-007, and the Department of Entomology at the University of California Riverside.

## Data Accessibility

The data reported in this study will be deposited in the Open Science Framework database upon acceptance, https://osf.io (DOI XXX).

## DISCLAIMER

Mention of trade names or commercial products in this publication is solely for the purpose of providing specific information and does not imply recommendation or endorsement by the U.S. Department of Agriculture. The conclusions in this report are those of the authors and do not necessarily represent the views of the USDA. USDA is an equal opportunity provider and employer.

## SUPPLEMENTAL FIGURES

**Figure S1.**
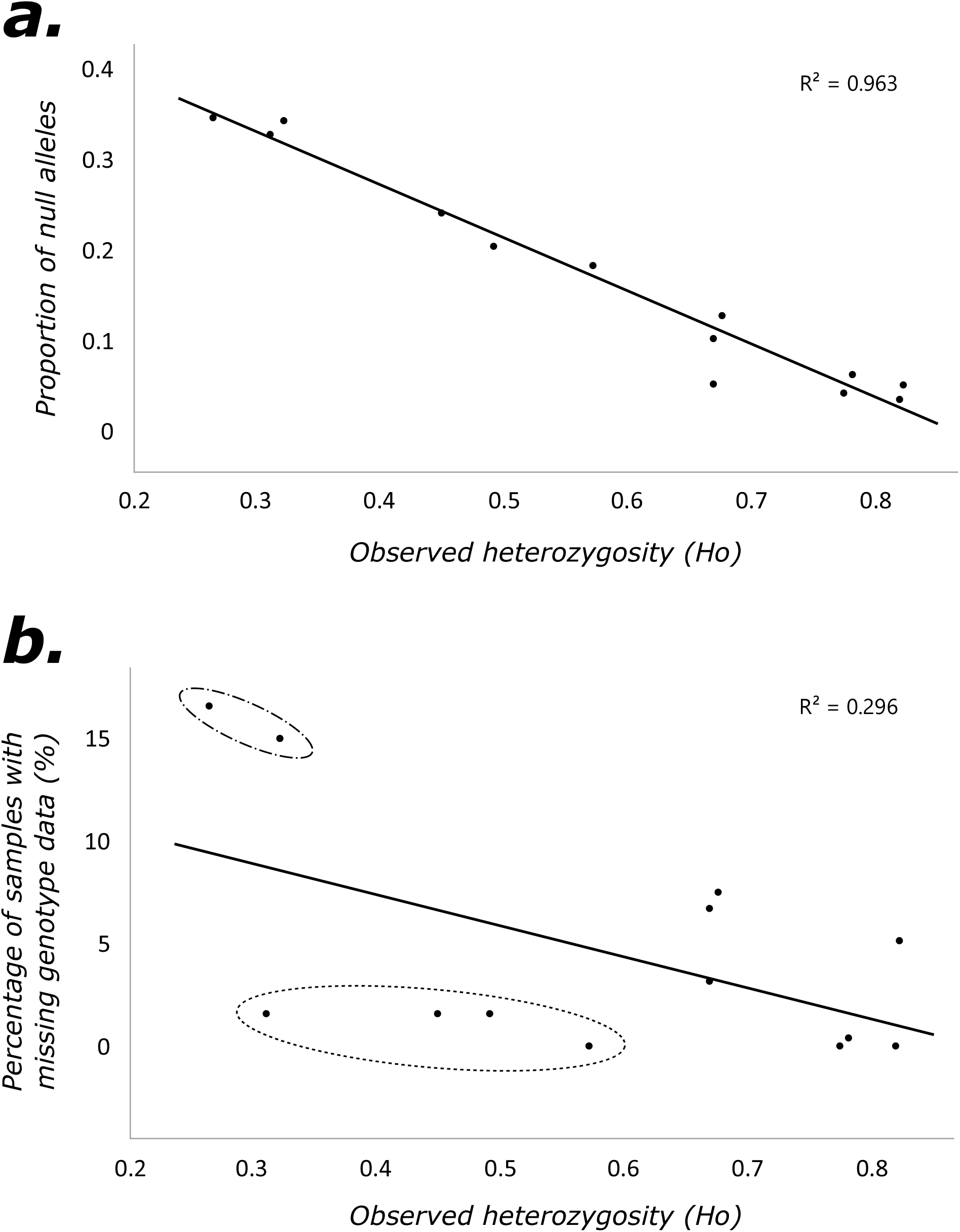
Estimates of the proportion of null alleles for each marker. (a.) A linear regression of the estimated proportion of null alleles and the observed heterozygosity. (b). A linear regression of the percentage of missing genotype data and the observed heterozygosity. The loci in the dashed/dotted circle have a high estimated proportion of null alleles and the supporting amount of missing genotype data. The loci in the dashed circle were also estimated to have a high proportion of null alleles; however, there is a discordant amount of missing data (< 2%).

**Figure S2.**
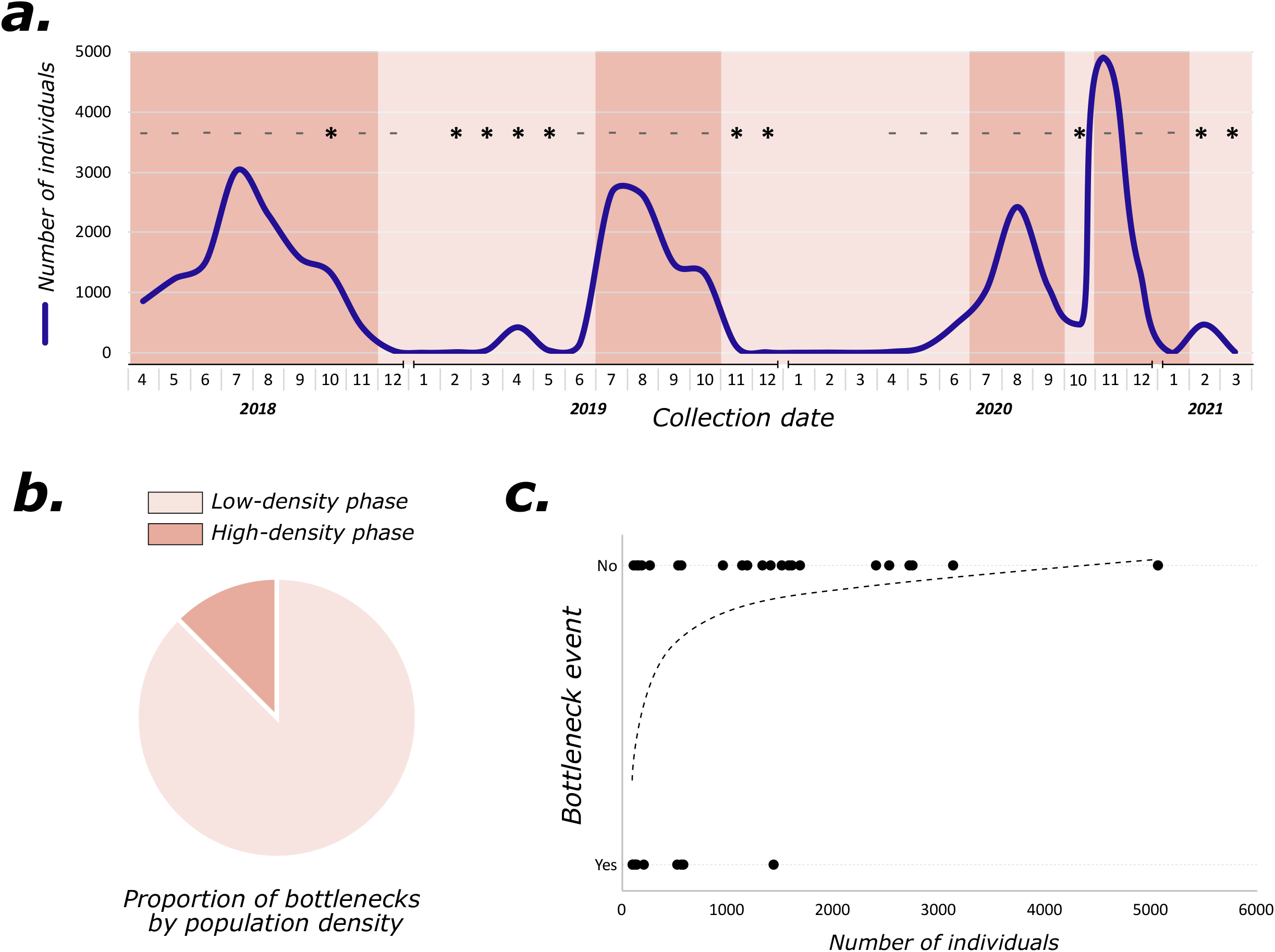
Inferred bottleneck events. (a.) The line shows the total number of individuals collected each month with the shaded area indicating either high (dark) or low (light) population densities. Above each month, a dash (−) denotes no bottleneck, an asterisk (*) denotes the inference of a bottleneck under the IAM, and no symbol indicates a month with no genetic information. (b.) The proportion of collection dates (N = 9) inferred to have a bottleneck event based on the corresponding adult population size. (c.) The relationship between adult population size and whether or not a bottleneck event was inferred.

